# Hallucinations both in and out of context: An Active Inference Account

**DOI:** 10.1101/540419

**Authors:** David Benrimoh, Thomas Parr, Rick A. Adams, Karl Friston

## Abstract

Hallucinations, including auditory verbal hallucinations (AVH), occur in both the healthy population and in psychotic conditions such as schizophrenia (often developing after a prodromal period). In addition, hallucinations can be in-context (they can be consistent with the environment, such as when one hallucinates the end of a sentence that has been repeated many times), or out-of-context (such as the bizarre hallucinations associated with schizophrenia). In previous work, we introduced a model of hallucinations as false (positive) inferences based on a (Markov decision process) formulation of active inference. In this work, we extend this model to include content – to disclose the computational mechanisms behind in- and out-of-context hallucinations. In active inference, sensory information is used to disambiguate alternative hypotheses about the causes of sensations. Sensory information is balanced against prior beliefs, and when this balance is tipped in the favor of prior beliefs, hallucinations can occur. We show that in-context hallucinations arise when (simulated) subjects cannot use sensory information to correct prior beliefs about hearing a voice, but beliefs about content (i.e. the sequential order of a sentence) remain accurate. When hallucinating subjects also have inaccurate beliefs about state transitions, out-of-context hallucinations occur; i.e. their hallucinated speech content is disordered. Furthermore, subjects with inaccurate beliefs about state transitions but an intact ability to use sensory information do not hallucinate and are reminiscent of prodromal patients. This work demonstrates the different computational mechanisms that may underlie the spectrum of hallucinatory experience – from the healthy population to psychotic states.

## INTRODUCTION

Hallucinations, including auditory verbal hallucinations (AVH), are a key component of the nosology and phenomenology of schizophrenia (APA, 2013). Despite decades of phenomenological, neurobiological and computational research, it is still not clear precisely how AVH develop and what mechanism underwrites them. Some investigators have turned to computational modelling to better understand the information processing deficits underlying AVH (Benrimoh et al., 2018; Powers et al., 2017; Jardri and Denève, 2013; Hoffman et al., 2006). Models based on Bayesian inference – inference via a combination of prior beliefs and sensory information (Benrimoh et al., 2018; Jardri and Denève, 2013; Powers et al., 2017) – support the notion that the overweighting of prior beliefs (e.g. that a certain word or sentence will be heard), relative to sensory information is necessary for hallucinations to emerge. We have shown that hallucinations emerge when an overweighting of prior beliefs is coupled with a decrease in the precision of, or confidence in, sensory information (Benrimoh et al., 2018). In other words, computational agents hallucinate when they believe they should be hearing something and are unable to use sensory information (e.g. silence) to update this belief. Empirical evidence supports this model. For example, using a hierarchical Bayesian model (the Hierarchical Gaussian Filter; Mathys et al. 2011) to fit data from a task in which participants were conditioned to hear certain tones, Powers et al. (2017) found that patients with hallucinations in the context of schizophrenia and healthy voice-hearers placed greater weight on their prior (i.e. conditioned) beliefs about the tone-checkerboard pairing. Teufel et al. (2015) found that individuals in early psychosis, and healthy individuals with psychosis-like experiences, had improved performance – due to greater contributions of prior knowledge – in a visual task compared with controls. Finally, Vercammen et al. (2010) found that healthy participants who were more hallucination-prone were more likely to hallucinate a predictable word (given the semantic context) when the stimulus presented was white noise or an unpredictable word partially masked by white noise (i.e. a stimulus of low precision).

This body of work, which has investigated the genesis of hallucinations resulting from an imbalance between prior beliefs and sensory information, is well-suited to describe ‘conditioned’ hallucinations (Powers et al., 2017), those hallucinations resulting from a prior that a subject acquires through recent experience, such as a pattern of tones. However, clinical hallucinations tend to have some additional properties: i) they are often ‘voices’ with semantic content and ii) they are often not coherent with the current environmental context. Semantic content can be derived from – or coloured by – a patient’s memory or feelings of fear (McCarthy-Jones and Fernyhough, 2011; McCarthy-Jones et al., 2014). Conditioned hallucinations are necessarily coherent with some aspect of the environmental context (it is this context from which the prior is inherited). Hallucinations in patients with schizophrenia are often emphatically *out of context*, however, such as a frightening voice that is heard in an otherwise safe room or hearing the voices of world leaders whom one has never met.

In this paper, we explore the computational mechanisms that could underwrite the difference between hallucinations that are *in*-context (like a conditioned hallucination) or *out-of-*context (closer to a true clinical hallucination). Our purpose in doing so is twofold. First, we aim to determine whether the same computational mechanism (Benrimoh et. al., 2018) is appropriate for describing hallucinations in and out of context. This is important for understanding the relevance of empirical work on conditioned hallucinations to the clinical phenomena. Our second aim is to understand, if the underlying mechanism is the same, what sort of additional computational deficits might be in play to give out-of-context hallucinatory percepts. We have previously argued that prior beliefs, when combined with reduced confidence in sensations, could underwrite hallucinations (operationalised as false positive inferences). In the present work, we investigated (through simulation) whether a similar mechanism could influence a subject’s beliefs – not just about whether they hear sound – but about the content of that sound. We hoped that the form of these hallucinations might enable us to draw parallels with known AVH phenomenology, and to show whether common mechanisms could underwrite both AVH and other psychotic symptoms, such as thought disorder.

## METHODS

To address the questions above, we simulated belief-updating in a simple *in silico* subject who could make inferences about the content she was hearing and could choose to speak or listen to another agent. To accomplish this, we used a Markov Decision Process (MDP) implementation of active inference (Parr and Friston, 2017; Mirza et al., 2016). We opted for an MDP as this furnishes a generative model of discrete states and time, where the recognition and generation of words are thought to occur in a discrete manner (Dietrich and Markman, 2003). Working within the active inference framework treats perception as an active process (i.e., active listening), in which an agent’s decisions (e.g. choosing to speak or listen) impact upon their perceptions (Parr and Friston, 2017). This is a crucial difference from previous models of AVH, which only consider passive perception (Adams et al., 2013; Hoffman et al., 2006). This is important because priors engendering hallucinations may concern action plans (or policies) an agent chooses to engage in, where these policies are ill-suited to the current context. For example, if the agent strongly believes it should be listening to another agent speaking, this prior is inappropriate if there is nothing to listen to. In this section, we describe active inference and the MDP formalism, and then specify the implementation of our *in-silico* agent.

### Active inference

Under active inference, creatures use a generative (internal) model to infer the causes of their sensory experiences and to act to gather evidence for their beliefs about causes. More formally, agents act to minimize their variational free energy (Friston et. al, 2006); a variational approximation to the negative log (marginal) likelihood of an observation, under an internal model of the world (Friston, 2012). This quantity – also know as surprise, surprisal and self-information – is negative Bayesian model evidence. This means that self-evidencing (or finding evidence for one’s model of the world) is the same as minimising variational free energy. The free energy can be written as:

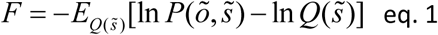

In this equation, *F* is the free energy, 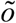 is the sequence of observations over time, *s* are unobserved or hidden states that cause sensations, and *Q* is an approximate probability distribution over *s*. *P* is the generative model that expresses beliefs about how sensory data are generated. This is the model that attempts to link observations with the hidden states that generate them.

### Markov Decision Process and Generative Model

MDPs allow us to model the beliefs of agents who must navigate environments in which they have control over some, but not all, variables. These agents must infer two types of *hidden* variables: *hidden states* and *policies*. Hidden variables are those variables that cannot be directly observed. The hidden states *s_τ_*, represent hypotheses about the causes of sensations; namely, current states of affairs generating the sensorium. For our agent, there are three sorts of hidden states: listening (or not); speaking (or not); and lexical content (i.e. the words that the agent may say or listen to). Intuitively, this kind of structure may be thought of as analogous to the factorisation of visual processing into ‘what’ and ‘where’ streams. Just as an object retains its identity no matter where it appears in space, the content of speech is the same regardless of which conversational participant is speaking.

This generative model is closely analogous to that used in previous treatments of pre-linguistic (continuous) auditory processing (Adams et al., 2013). These simulations relied upon generation and perception of synthetic birdsong. Crucially, this birdsong had a pre-defined trajectory that depends upon internally generated dynamics. Similarly, our simulations depend upon an internal conversational trajectory; i.e., narrative. This comprises the six words in the short song “It’s a small world after all!” but does not specify which word is spoken by which agent. The hidden states are inferred from the sensory observations *o_τ_* (where *τ* indicates the time-step). Our agent has access to two outcome modalities: audition and proprioception. The auditory outcome includes each of the six words in the song, as well as a seventh ‘silent’ condition, which is generated when the agent is neither listening or speaking.

As previously (Benrimoh et al., 2018), our subject’s generative model is equipped with beliefs about the probability of a sensory observation, given a hidden state (i.e. the probability of observing some data, given a state). These probabilities can be expressed as a likelihood matrix with elements *P*(*o_τ_* = *i* | *s_τ_* = *j*) = A_*ij*_. For the proprioceptive modality, this was simply an identity matrix (mapping speech to its proprioceptive consequences). For the auditory modality the likelihood depended on whether the agent believed it was listening or speaking (in which case it did not consider silence to be possible and had non-zero probabilities for the six content states, see Figure 1) or if it believed no sound was present (in which case it assigned a probability of 1 to the silent outcome). The resulting likelihood matrix was then passed through the following equation:

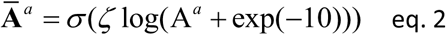

**Figure 1.**
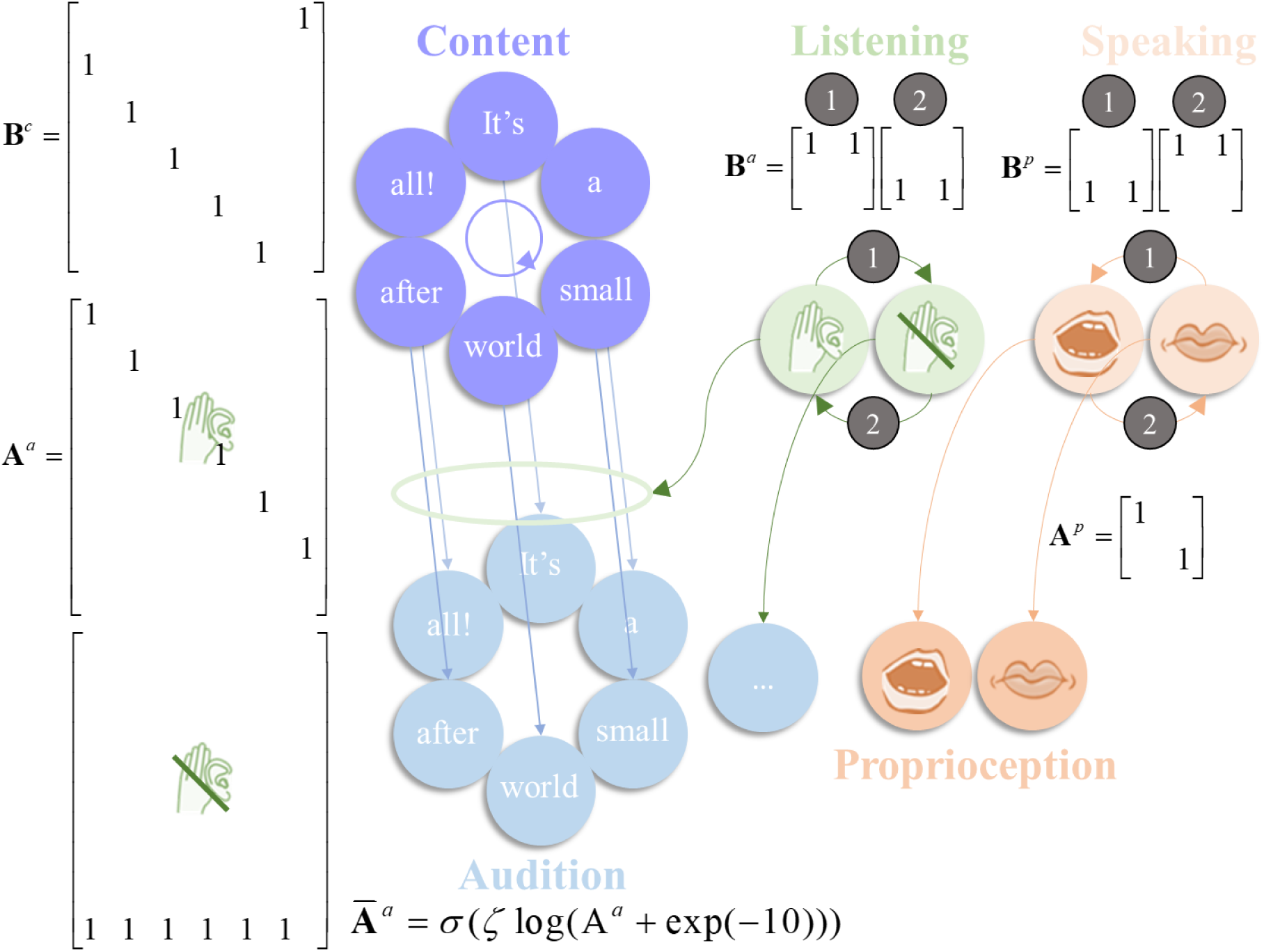
The generative model. This schematic illustrates the specification of the (partially observed Markov decision process) generative model used in the simulations. There are three sorts of hidden state: those that determine the content of speech (i.e. the words in the song), whether the agent is currently listening to anything, and whether the agent is speaking. The first deterministically transitions from each word to the next in the song, and repeats this sequence indefinitely as shown in the **B**-matrix for this content state Β^*c*^. The latter two hidden state factors depend upon the agent’s choice; with actions given in small numbered circles. The policies are defined such that, when action 1 (Speaking) is taken for the listening state (leading to ‘I am not listening’), action 1 is also taken for the speaking state (leading to ‘I am speaking’), and similarly for action 2 (Listening). There is no such constraint in the generative process, so this belief can be violated. When listening, the (lexical) content states map to auditory outcomes representing the same words. This is shown in the first of the auditory **A**-matrices A^*a*^ for which columns represent content states, and rows content outcomes. The second (lower) version of this matrix is employed when not listening to anything and gives rise to a ‘silent’ auditory outcome (7^th^ row). There is additionally a proprioceptive outcome modality *o*^*p*^ that depends only upon whether the agent is speaking. The equation at the bottom of this Figure shows how we equip the auditory likelihood with a precision (inverse temperature) parameter *ζ*. The bar above the A indicates that the matrix has been normalised, following precision modulation.

Here, *σ* is the softmax function which normalizes the matrix. *ζ* is the likelihood precision^1^. The bar notation means that **A**^*a*^ has been normalised (the superscript denotes the auditory modality). As *ζ* increases, this mapping tends towards an identity matrix (that is, the mapping between causes and sensations becomes more deterministic). As it decreases, the probabilities become close to uniform and the mapping becomes very uncertain: i.e., even if one knew the hidden state of the world, all outcomes are equally likely. In other words, modulating the likelihood precision modifies the fidelity of the mapping between states and outcomes; the lower the precision, the ‘noisier’, or stochastic, the mapping becomes. In continuous state space formulations of active inference, optimising the equivalent quantity is synonymous with the process of directing attention; by attending to a sensory channel, an agent increases the precision of information it receives from that channel (Feldman and Friston, 2010). We have previously shown that synthetic agents tend to ‘disregard’ low-precision mappings (Parr & Friston, 2017) (c.f. the ‘Streetlight effect’ (Demirdjian et. al, (2005), which states that the sensible place to look for information is where it is most precise; i.e. under a streetlamp in an otherwise dark street).

Policies are the second type of hidden variable. Policies are sequences of actions (e.g., listening or speaking) that an agent can employ. Crucially, the currently pursued action must be *inferred* from sensory data. This is known as planning as inference (Botvinick & Toussaint, 2012). For our agent, policies are sequences of actions that generate a conversational exchange. Specifically, our agent has three allowable policies: always listen, always speak, or alternate between speaking and listening. At any given time, the policies are mutually exclusive. In other words, our subject believes she is listening or speaking, but not doing both at the same time.

The hidden states change over time according to a probability transition matrix, conditioned upon the previous state and the current policy, *π*. This transition matrix is defined as Β(*u*)_*ij*_ = *P*(*s*_*τ*+1_ = *i* | *s*_*τ*_ = *j*, *u* = *π* (*τ*)). The policy influences the probability a state at a given time transitions to a new state (e.g. if I choose to listen, I am more likely to end up in the ‘listening’ state at the next time-step). Heuristically, we consider ‘listening’ as an action to be a composite of mental and physical actions – all the things one might do to prepare to listen, when expecting to hear someone speak (i.e., pay attention, turn your head to hear better, etc.) (Holzman, 1972). While listening and speaking are actions and are therefore selected as a function of the inferred policy, the lexical content of the song from beginning to end is not. In other words, the order of the words in the song – i.e. the transitions between different content states – is a prior stored in Β^*c*^, the probability transitions encoding some level or latent narrative (e.g., song). The associated transition matrix could take one of two forms – as can be seen in Figure 1: a *veridical* form, where the song progresses in a standard fashion, starting at “It’s” and ending at “all!”; alternatively, *anomalous* narrative could be in play, which provides a ‘private’ narrative that is not shared with other people. This alternative perceptual hypothesis will form the basis of the anomalous inference we associate with hallucinations. It is important to note that the specific anomalies induced in these simulations are not intended as a literal account of schizophrenia; rather, the point we attempt to make is that an account of psychotic hallucinations requires a disruption of internally generated dynamics to account for the often-bizarre narratives, characteristic of such hallucinations. This does not necessarily influence whether a hallucination takes place, but profoundly alters what is hallucinated.

The subject was paired with another agent (i.e., a generative process) that determined the sensory input experienced – a partner that produces alternating sounds and silences according (always) to the normal content of the song (i.e. it produces every second word, beginning with ‘It’s’). Whenever our subject chose to speak, a sound was generated. Our subject could also choose to listen to – or sing with – her conversational partner. In this way, our subject interacted with the (simple) generative process, generating sequences of sounds and silences depending on both the environment and the actions. We chose a dialogic set up to provide the subject with context, which allows us to demonstrate out-of-context hallucinations. In addition, having a dialogic structure allowed us to more closely mimic hallucinatory phenomena, which can take the form of dialogue (McCarthy-Jones and Fernyhough, 2011).

The subject began a prior probability distribution over possible initial hidden states, D_*i*_ = *P*(*s*_1_ = *i*). In our simulations, our subject always listened at the first timestep, as if she was ‘getting her cue’. The subject was also equipped with a probability distribution over possible outcomes, which sets the *prior preferences*, C_*τi*_ = *P*(*o_τ_* = *i*). These prior preferences influence policy choices, because agents select policies that are likely to lead to those outcomes. Here, we used flat priors over outcomes, ensuring no outcome was preferred relative to any other. This means that action selection (i.e. talking and listening) was driven purely by their epistemic affordance, or the imperative to resolve uncertainty about the world.

To make active inference tractable, we adopt a mean-field approximation (Friston and Buzsaki 2016):

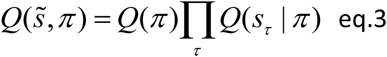

Here *Q* represents approximate posterior beliefs about hidden variables. This formulation (which just says that the approximate posterior beliefs about hidden variables depends on the product of the approximate probability over policies and the approximate probability of states given polices at a given time) allows for independent optimization of each factor on the right-hand side of eq. 3. The expression of the free energy under a given policy is:

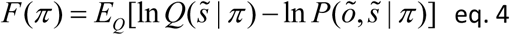

Free energy scores the evidence afforded to a model by an observation. This poses a problem when selecting policies, as these should be chosen to reduce future – not current or past – free energy (and we do not yet have access to future observations). In this setting, policies are treated as alternative models, and are chosen via Bayesian model selection, where the policy selected is the one that engenders the lowest expected free energy

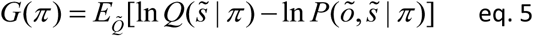

Here *G*(*π*) is the *expected* free energy under a given policy and allows for predictions about what is likely to happen if one pursues a given policy – that is, how much uncertainty will be resolved under the policy (and when prior preferences are used, how likely the policy is to fulfil them).

Formally, the distribution over policies can be expressed as

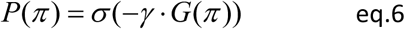

This equation demonstrates how policy selection depends on the minimization of expected free energy. This is equivalent to saying that agents choose policies that are most likely to resolve uncertainty. Here, the expected free energy under alternative policies is multiplied by a scalar and normalized via a softmax function. The scalar plays the role of an inverse temperature parameter that signifies the precision, or *confidence*, the agent has about its beliefs about policies. This confidence will have an impact on the relative weighting of sensory information, when the agent tries to infer its policy. The more confident an agent is about its policies, the less it relies upon sensory information to infer its course of action. The *γ* parameter has previously been linked to the actions of midbrain dopamine (Schwartenbeck et al., 2015). We have previously shown that an increased prior precision over policies can facilitate hallucinations (Benrimoh et al., 2018), but in this paper we do not modulate this policy precision.

In the above, posterior beliefs about states are conditioned upon the policy pursued, which is consistent with the interpretation of planning as model selection. Through Bayesian model averaging, we can marginalize out this dependence to recover a belief about hidden states, irrespective of beliefs about the policy pursued:

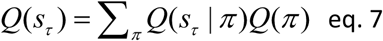

This equation is crucial because it means that beliefs about states (to the left of the equation) depend upon beliefs about the policy. The terms on the right correspond to the approximate probability distribution over policies, and over states, given policies. This fundamental observation further ties perception to action – and frames the development of hallucinations as an enactive process, which was the focus of our previous work (Benrimoh et al., 2018). In that work, we showed that policy spaces – prior beliefs about action sequences – which did not match sequences of events in the environment led to hallucinations when external information could not be used to correct perceptions (because of low likelihood precision). In this paper, we use anomalous policies to characterise perceptual inference (about lexical content) during hallucinations.

### Hypotheses: The Generation of In-Context and Out-of-Context Hallucinations

To summarise, we have an agent with policies for active listening, who is tasked with singing a duet with another agent. Our agent can be equipped with either a veridical narrative (i.e., transition matrix); namely, the belief that the song will always proceed as “It’s a small world after all!”. Alternatively, it can be equipped with an anomalous transition matrix –under which it believes that the words should be sung in a different order. Crucially, the presence of the other agent – who always sings the song in the standard order – provides a *context* for our agent. Our objective was to produce in and out-of-context hallucinations, and to see which factors disambiguate between the two.

From our previous work, we hypothesized that our agent would hallucinate when the likelihood precision was reduced relative to prior beliefs. We further hypothesized that, when this occurred with a veridical transition matrix, the agent would hallucinate perceptual content *consistent* with the context. This is because the agent’s beliefs about lexical content are entirely consistent with the narrative being followed in the environment. In contrast, we hypothesized that the agent would hallucinate *out-of-context*, when equipped with the anomalous transition matrix. This is because when the likelihood precision is low, our agent is unable to use sensory information to correct its anomalous beliefs regarding lexical transitions. For the same reason, we expected the agent with the anomalous transition matrix to infer the standard word order when likelihood (i.e., sensory) precision was high – because it would be able to use the information present in the environment to correct its biased prior beliefs about content state transitions. In short, with this relatively straightforward setup, we hoped to demonstrate a dissociation in terms of context sensitive hallucinations, and active listening

## RESULTS

Here, we present the results of our simulations. These are presented in Figures 2-5 and represent the results of four *in silico ‘*experiments’. In Experiment 1, we showcase a subject that demonstrates normal behavior and non-hallucinatory beliefs. This subject is equipped with a veridical transition matrix and as such believes that the song will unfold in the usual way. This is reflected in the purple ring of circles (and its adjacent matrix) in Figure 1. The agent is also equipped with a normal likelihood precision, and as such can accurately use sensory information to infer hidden states. For example, it accurately infers the silent state at the second and fourth timesteps. At other times, it chooses to sing or listen to the other agent singing, but in all cases its beliefs about the lexical narrative map perfectly onto auditory outcomes.

**Figure 2:**
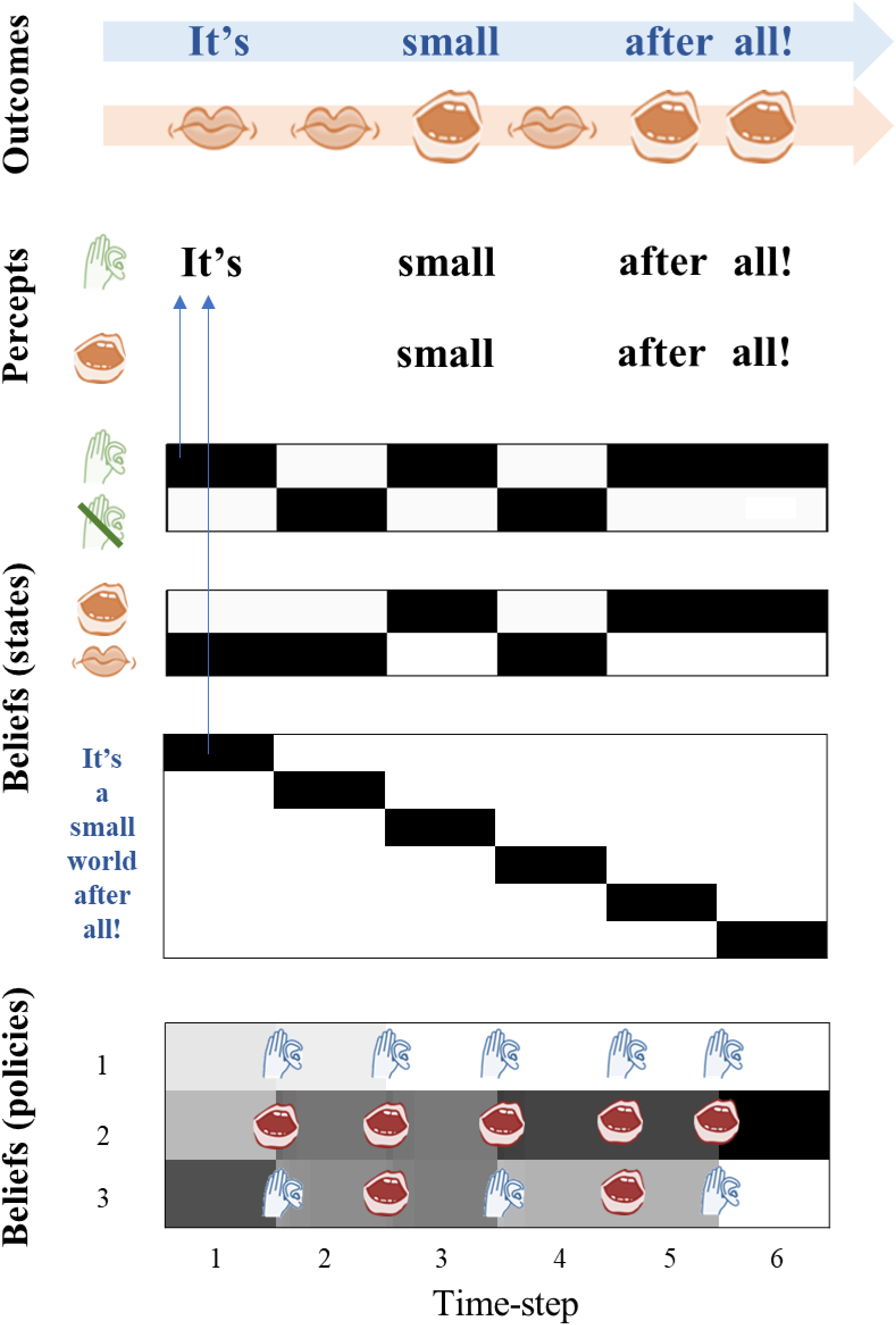
Normal State. This figure depicts the normal state of the agent, without any induced pathology. These are the results of the simulation for six timesteps. True outcomes are listed at the top. The agent’s posterior beliefs about states (audition, proprioception, and content) as well as its beliefs about lexical content are shown. The three policies are depicted in the lower half of the figure, with each possible action situated at the end of a timestep – one of which is sampled probabilistically as time transitions to the next timestep. N.B. the actual action sampled is not shown; neither are the actions of the confederate agent. For all figure elements with black and white shading, black denotes high certainty; white denotes low certainty; shades of grey denote intermediate certainty. In this figure the agent hears words in the order of the song and does not experience any hallucinations. The sequence of events is as follows: Timestep 1 – The agent is ‘cued’ – it expects to be listening to the other agent and not speaking in the first time-step. This initial state is common to all the simulations. Timestep 2 – The agent samples an action from the most likely policy given the previous timestep (3), which is Listening. However, it hears only silence (because the other agent sings only on timesteps 1, 3 and 5), so it infers its state is ‘I am not listening’. It is not speaking so it also infers ‘I am not speaking’. Given these states, its posterior beliefs about its policy are equivocal between the listening and speaking policies (all are unlikely). Timestep 3 – The agent samples an action from policy 2 or 3: both are Speaking. It speaks and infers ‘I am speaking’ because its mouth is moving. It also hears a sound, so it infers ‘I am listening’ and the content it infers to be ‘small’ because it is up to this point in the transition matrix. Timestep 4 – At the start of this timestep it is equivocal between policies 2 and 3 as both were speaking at the last timestep. It samples an action – Listen – so its mouth does not move, and it correctly infers ‘I am not speaking’. However, it hears only silence, so it also infers its state is ‘I am not listening’. Timesteps 5 & 6 – The agent samples from the most probable policy (2): it action is Speaking. It speaks, its mouth moves, and so it infers its state to be ‘I am speaking’. It also hears sounds at each time point which make it infer ‘I am listening’ and the content to be the 5^th^ and 6^th^ entries in the content transition matrix.

**Figure 3:**
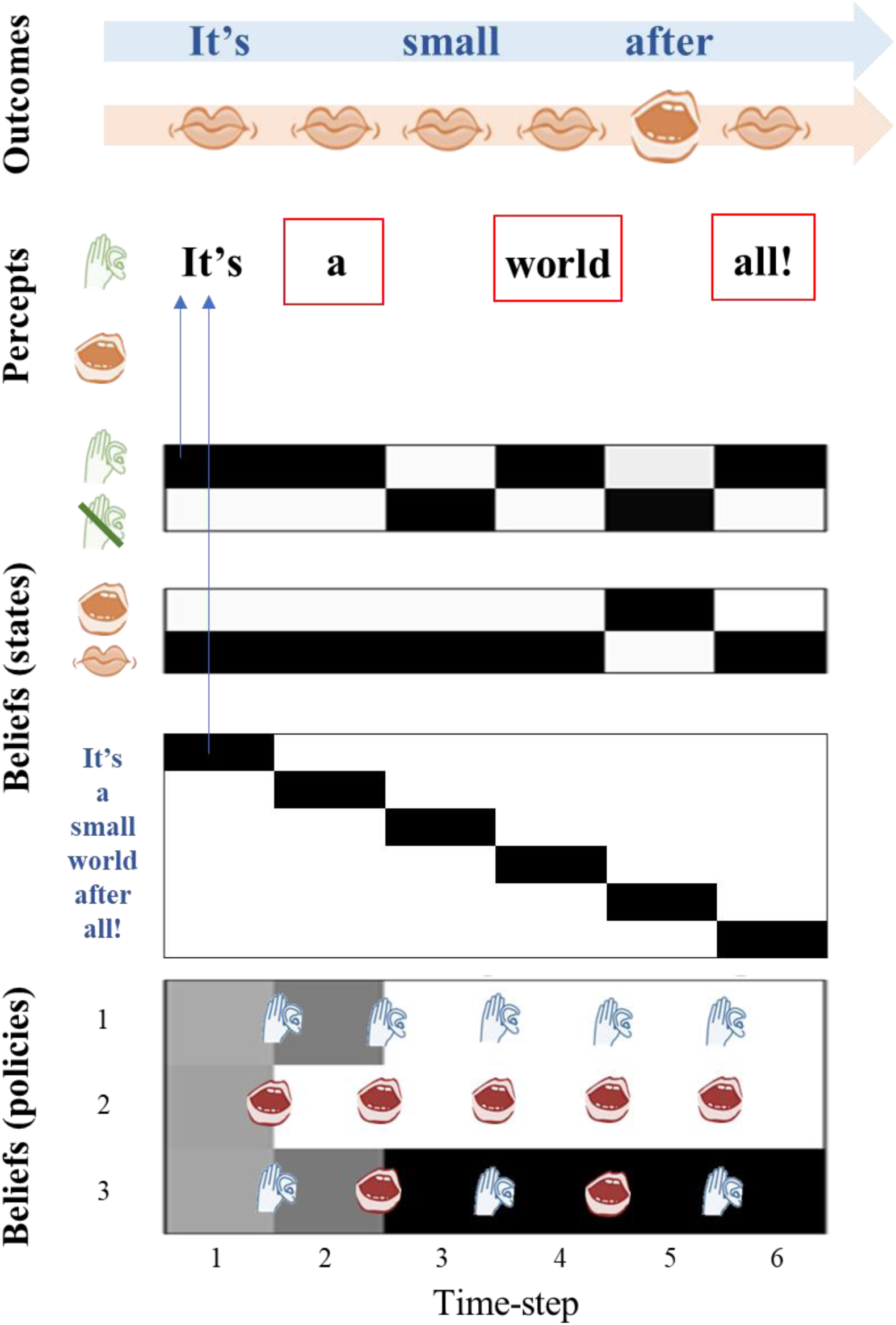
In-context Hallucinations. In this figure, the agent has been given a reduced likelihood precision but has access to the same policies and the same transition matrices as the agent in Figure 2 (the ‘normal state’ agent). With this lowered precision, the agent hallucinates at the 2^nd^, 4^th^, and 6^th^ time steps (red boxes), hearing the words of the song it expects to be present at those time steps based on its transition matrix. It also fails to hear its own voice at the 5^th^ time step, and the voice of the other agent at the 3^rd^ time step. At time-step 1, the agent expects to be ‘cued’. Auditory input is present, but there is no proprioception, and so the agent correctly infers it is listening but not speaking; all the policies are roughly equally probable at this point. At the second time-step, there is no audition or proprioception and the agent infers that it is not speaking (due to high proprioceptive precision). This is consistent with policy 1 or 3 having been engaged at the previous time-step, but inconsistent with policy 2. As the agent infers policies 1 and 3 with high confidence, this provides a strong prior belief that it must be listening during this time-step. As there is low likelihood precision, the agent hallucinates because it cannot use the information in the environment (i.e. the absence of sound) in order to correct this prior belief. At the 3^rd^ time-step, there is auditory information but no proprioception; the agent should infer that it is listening but not speaking, which would be consistent with both modalities. However, the low likelihood precision leads the agent to fail to hear the sound and the agent infers it is not listening. This is inconsistent with policy 1, so this is eliminated, and the agent becomes confident in policy 3, which it then employs for the rest of the simulation. This means that at the 4^th^ and 6^th^ timesteps, the agent is very confident it is listening, but its low likelihood precision means it cannot accurately perceive the absence of sound, so it hallucinates – just as in timestep 2.

**Figure 4:**
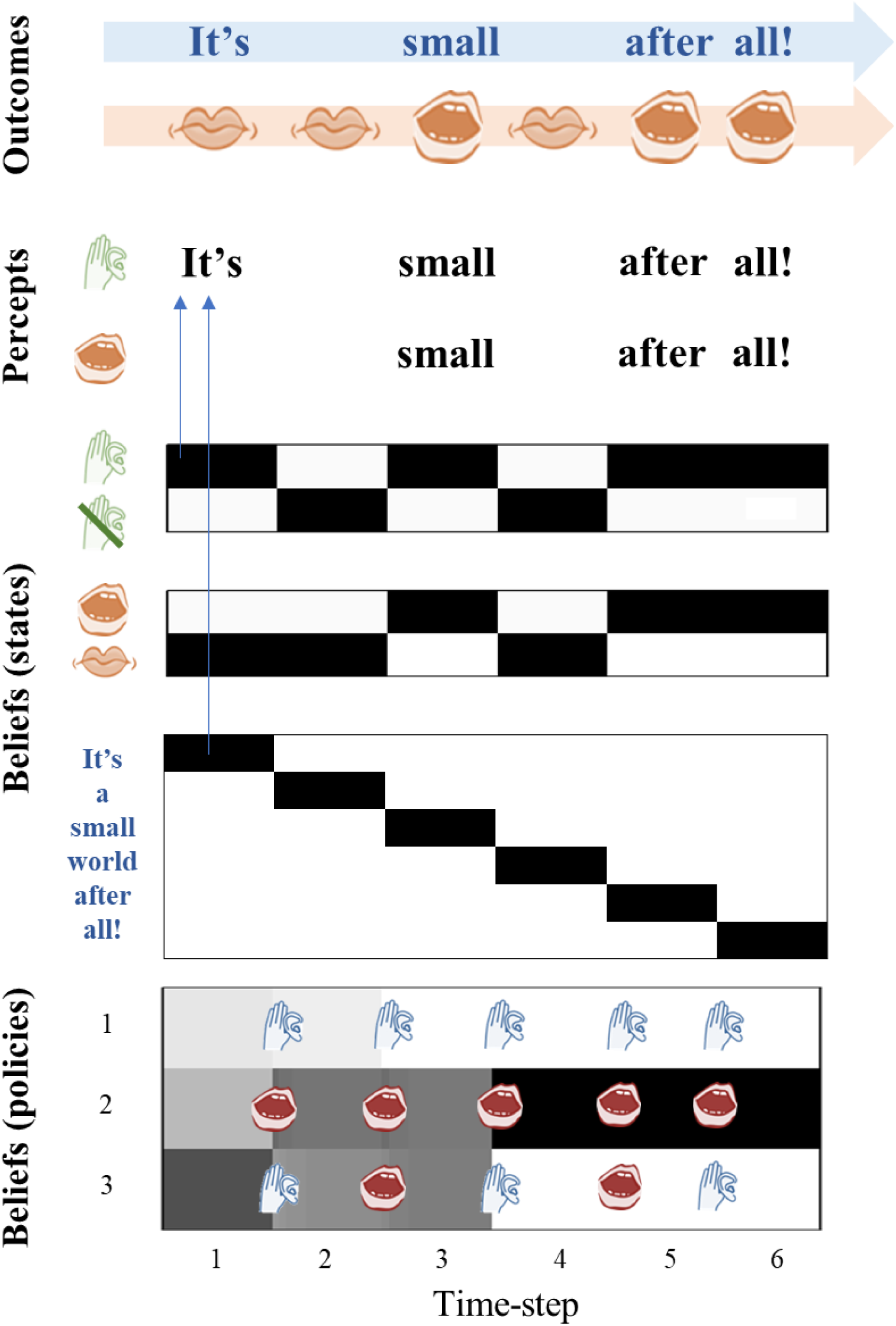
“Prodromal” Agent. In this figure, the agent has been equipped with an anomalous transition matrix for the lexical state, but it retains a normal likelihood precision. This enables the agent to avoid any hallucinations, and it perceives the order of words expected given the order of the song as sung by the generative process (i.e. the normal song order), even though it has been biased to expect a different transition structure. This can be considered a ‘prodromal’ agent because it has anomalous beliefs that are kept from being expressed or experienced as hallucinations because of the agent’s ability to use sensory information to correct its beliefs. Note that at the end of the third timestep, the agent has equal probabilities for policies 2 and 3 and could sample either a speaking or listening action. At the fourth timestep, there is no sound as the agent does not sample a speaking action and the other agent does not speak; as the agent has a normal likelihood precision it is able to correctly infer silence.

**Figure 5:**
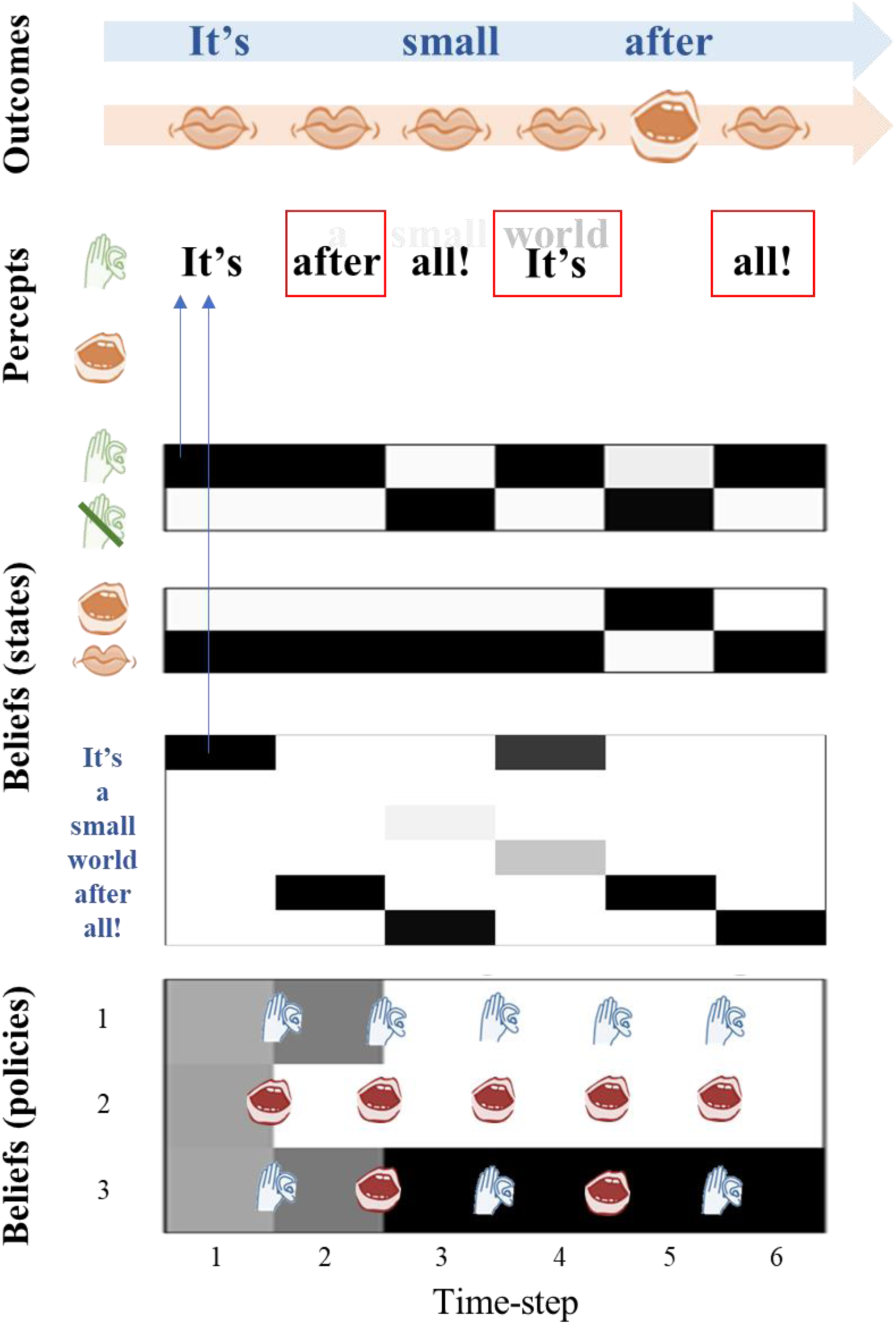
Out-of-context Hallucinations. In this figure, the agent has been equipped with an anomalous transition matrix as well as a reduced likelihood precision. In this case, the agent again experiences hallucinations at the 2^nd^, 4^th^, and 6^th^ time steps but these are ‘out of context’ or out of the usual order of the song at the 2^nd^ and 4^th^ timesteps. Here the agent has heard “after” at the second time step instead of “a”, and “It’s” at the 4^th^ time step instead of “world”. It also mishears the other agent (the generative process) at the 3^rd^ time step, hearing “all!” instead of “small”. It fails to hear its own voice at the 5^th^ timestep. Here, lighter colored font indicates a lower certainty that word is being heard, and darker font a higher certainty. As an example of how an out-of-context hallucination emerge, going into the 2^nd^ timestep the agent samples a “listening” action, from policy 1 or 3. This induces a prior belief that it will be hearing something at timestep 2, even though no sound is present at this time. The low likelihood precision leads it to hallucinate – as it cannot use the silence to correct this prior belief. As the anomalous transition matrix is in play, the agent hears “after” instead of “a”, because it is unable to use the lexical cues to predict the correct word order.

In Experiment 2 we reduced the likelihood precision (from *ζ* = 3 to *ζ* = 1), which engendered hallucinations at the second, fourth and sixth timesteps. As discussed in our previous paper (Benrimoh et al., 2018), these hallucinations are driven by priors derived from the policies selected by the agent: these were all timesteps immediately before which the ‘listening’ action had been selected, and as such the agent expected to be listening (i.e., attending) to sounds. Given reduced confidence in its sensations, the agent disregarded its observations (of silence) and instead inferred that sound was present, as expected. What is critical here is that *what* it heard was determined by its beliefs about lexical content and word order. As such, all the hallucinations were in-context, in the sense that they reproduced the sound as expected in the sequence present in the environment. At the third timestep, this subject failed to hear her partner, and at the fifth timestep she failed to hear her own voice. This is again a symptom of selecting a policy ill-suited to the context: i.e. one that did not cohere with states of affairs in the environment. At the third timestep, the subject is unsure of whether to listen or speak (follow policy one or two), and at the fifth timestep it has selected the third policy – which leads it to choose to speak and to believe it will not be listening. However, under normal likelihood precision the agent would be able to use sensory information to correct that belief; i.e. to infer it was listening, even if it was speaking or did not expect to hear anything – as seen in Experiment 1, when the subject hears her own voice in the last timestep, even though she was speaking). It should be noted that Hoffman et al., (2006) found that misperception – including the failure to perceive certain sounds – is a feature of hallucinations.

In Experiment 3, we equipped our subject with an anomalous transition matrix and a normal likelihood precision. If the subject’s beliefs about lexical content were solely determined by its priors, we would expect her to believe that the song was being sung in an anomalous order. However, because it is coupled to an environment that provides context, it can use this context to correct its priors and infer that the song is, in fact, following its usual order.

In Experiment 4, we removed the ability to correct anomalous perception by reducing the likelihood precision, which leads to the development of hallucinations and to an anomalous perceptual experience. The subject now believes that instead of the song progressing as “It’s a small world after all!” She perceives “It’s after all! It’s after all!”. Note that the policy selection – and therefore the priors that drive the hallucinations – are the same as in Experiments 2 and 4. The only thing that has changed are the beliefs about lexical (i.e., hidden state) transitions.

## DISCUSSION

In this paper, we simulate an agent who can experience both in-and out-of-context hallucinations. Reproducing our previous work, we show that hallucinations emerge when likelihood precision is low relative to prior beliefs. This induces false positive inferences that are consistent with the perceptual content expected in the presence of a perceptual narrative (e.g., dyadic singing or conversation). This may be thought of as analogous to the sorts of hallucinations induced in (often neurotypical) subjects in empirical studies. To investigate whether this mechanism generalises to the sorts of hallucinations associated with the clinical phenomenology of psychotic hallucinations, we asked what minimal computational pathology is required for the genesis of out-of-context hallucinations. Induction of imprecise prior beliefs about the trajectory of lexical content was enough to reproduce this phenomenon. This conceptual and computational analysis endorses the relevance of empirical studies that attempt to induce in-context hallucinations to understanding the basic mechanisms of hallucinogenesis in psychopathology, but additionally implicates abnormalities in internally generated narratives in understanding what is hallucinated.

The policy selections that drove hallucinations in Experiments 2 and 4 were identical. The difference between the two cases is specifically *what* the subject hallucinated, not *how* the hallucinations have come about. This dissociation between hallucination onset and hallucination content illustrates the differences between in-context and out-of-context hallucinations. The former lead to hallucinations whose content is governed by appropriate context (i.e., it matches the environmental context), whereas the latter are governed by priors that are not apt for a normal dyadic exchange. It is also interesting to note that the agent continues to attempt to use environmental cues to correct its beliefs about content, even under low likelihood precision. As can be seen in Figure 5, in the third and fourth timesteps the subject still has a weak belief that she is hearing the sound that would be encountered under a veridical song order. This may represent the uncertainty that some patients feel about the reality of their hallucinations (McCarthy-Jones 2014).

What is common to these results are that *out-of-context* hallucinations occur when an agent has a poor model of reality (i.e., anomalous priors) and dysfunctional method for checking the predictions of that model (relatively low likelihood precision). This is in keeping with a long tradition of understanding the psychotic state as a loss of contact with reality (Bentall et al., 1991). It is also reminiscent of Hoffman’s computational model (Hoffman et al., 2006). In that model, hallucinations were produced in the presence of degraded phonetic input (analogous to our imprecise likelihoods) and an overly pruned working memory layer, which allowed anomalous sentence structures to become possible (analogous to our anomalous transition matrix). By simulating a subject that can hallucinate content outside of the appropriate context, we are tapping into the disconnection from reality – and sensory evidence – that is central to schizophrenia, where patients can have sensory experiences that do not cohere with environmental context (hallucinations) and experience beliefs that are not supported by evidence (delusions). These beliefs may be strange and very much out of the realm of normal experiential context (bizarre delusions, such as those about extraterrestrials). The fact that these experiences feel very real to patients (McCarthy-Jones et al., 2014), while being very much out of the context provided by every day experience, speaks to a mechanism that allows for both a detachment from – and an altered model of – reality, such as the one we propose involving reduced sensory (i.e., likelihood) precision.

It is interesting to consider the lexical content perceived in Experiment 4. The subject hears a ‘nonsense’ version of the song, one in which the correct order of the words has been lost and repetition sets in. While this is a result of the set-up of this simulation – which makes it possible for the agent to experience only this song or silence – it is interesting to note that 21% of patients with hallucinations do in fact experience nonsense hallucinations (McCarthy-Jones et al., 2014). In addition, this out-of-order song bears a striking resemblance to another important feature of psychosis: thought disorder. Thought disorder can include thought blocking, loosening of associations between thoughts, repeating thoughts and perseveration, and extreme disorganization of thought (such as the classical ‘word salad’) (Andreasen, 1986). Our synthetic subject appears to experience thought disorder in the form of repeating nonsense phrases, caused by anomalous prior beliefs about lexical transitions, in conjunction with a reduced likelihood precision.

While thought disorder and hallucinations cluster separately in some symptom-cluster analyses (Arndt et al., 1991), our aberrant transition matrix only determines the *type* of hallucination, and not their presence or absence. As such, our simulations do not contravene the results of these cluster analyses. Furthermore, some theoretical models of hallucination draw links between cognitive dysfunction and thought disorder in the prodromal phase and later development of hallucinations, which may be influenced by these disordered thoughts. For example, as reviewed by Handest et al., 2016, Klosterkotter (1992) noted that many patients described elements of thought disorder in the prodromal phase and hypothesized that these could transform into audible thoughts. Conrad (reviewed in Sass and Parnas 2003) described “trema”, a state of fear and anxiety as well as cognitive and thought disturbances during the prodromal period, which is succeeded by frank psychosis and the development of hallucinations and delusions; he conceptualized certain thought disturbances, such as pressured thinking or thought interference, as transforming into hallucinations.

Our simulations do not include any affective or interoceptive aspects and, as such, we are not able to make any claims regarding the ‘mood’ or ‘anxiety’ the subject could be experiencing (i.e. we cannot say if it is experiencing trema). However, it is interesting to consider the clinical analogies from Experiment 3 in light of Klosterkotter’s and Conrad’s models. This agent would be experiencing some odd internal beliefs that must be reconciled with external reality (the environmental context), but it is still able to do this successfully and make correct state inferences. In Experiment 4, when the likelihood precision is decreased, we see an ‘unmasking’ effect, akin to the onset of psychosis: hallucinations occur, and the out-of-context beliefs that had been suppressed now manifest as out-of-context hallucinations. While this remains speculative, one might conceptualize the subject from Experiment 3 as being in the prodromal phase; namely, a phase that is possessed of odd, context-incongruent prior beliefs but which is able to maintain normal inference. When this subject loses her ability to correct odd prior beliefs by reference to the environment, she enters a true psychotic state – not only hallucinating but doing so in an out-of-context manner.

These simulations allow us to hypothesize about the neurobiological mechanisms that could underlie these computational states. An altered generative model of the world; e.g., containing imprecise or improbable (in the real world) transitions, could arise due to genetic and/or neurodevelopmental changes in synaptic and circuit function. Yet these changes may only manifest in abnormal inference, once likelihood precision is compromised; for example, in a disordered hyperdopaminergic state. A key question is therefore: what might cause the onset of such a state? There may be purely neurobiological reasons for this transition. Another possibility is that this change is at least in part computationally-motivated; i.e., Bayes-optimal, given some (pathological) constraints. In some circumstances, it may be optimal for an agent to reduce its sensory (likelihood) precision-for example, if sensory evidence conflicts with a prior belief held with great precision (see Parr et al. 2018 for a related simulation). The latter may be the result of powerful effects (e.g. an overwhelming sense of paranoia or grandiosity) or parasitical attractors in dysfunctional cortical networks (Rolls et al., 2008).

Therefore, a person beginning to experience thought disorder might at first try to use information from the environment to check or update her beliefs. Should these thoughts remain impervious updating by contradictory data – for the pathological reasons suggested above – the person might begin to decrease the precision afforded to external information and adopt a new transition structure. These aberrant priors may manifest as thought disorder, eventually leading to frank psychosis whose content is determined by this new transition structure and potentially themed according to the prevailing affect. As noted in McCarthy-Jones et. al., (2014), memories and fearful emotions are frequently the content of hallucinations, which would seem to implicate medial temporal structures, such as the hippocampus and amygdala, and the amygdala-hippocampal complex has also been shown to be abnormal in schizophrenia (Lawrie et al., 2003). Indeed, in those with visual hallucinations and schizophrenia, hyperconnectivity of the amygdala and visual cortex has been observed (Ford et al., 2015). While our model does not explicitly include memory or emotional states, future work could simulate belief updating that shapes hallucination content through the impact of memories or fear. The aim of this work would be to sketch a computational account of the prodromal phase and its subsequent development into frank psychosis.

## LIMITATIONS

Our model has several limitations. The first is that assumes a dialogic context, where the environment takes the form of another agent singing at every second time-step. This singing provide evidence for the environmental context. This context is rather simplistic and contrived, but it is intended to provide the a minimalist illustration of contextual and non-contextual hallucinations. It is not clear if agents would respond the same way to other forms of environmental context, such as visual cues, and if these would serve as the context for verbal hallucinations. In addition, we selected certain types of priors – those pertaining to action selection and state transitions – and it should be clear that these are not the only possible sources of priors for hallucinations. For example, an agent might simply have a strong preference for hearing certain content (e.g., caused by strong affective states), which could bias policy selection towards hearing that content, but this is not something we have explored in this simulation.

For simplicity, and to reinforce the dialogic structure of the simulation, we restricted our policy spaces so that speaking, and listening were mutually exclusive during action selection. However, the purpose of the simulation was not to comment on specific sequences of inferred actions that may exist *in vivo*, and as such this set-up does not impact upon the meaning of our simulations. In addition, we only employed two modalities – proprioception and audition – and we assumed no errors in proprioception, given the absence of gross sensory deficits in schizophrenia.

We also omitted a key phenomenon in schizophrenia from our model: the failure of sensory attenuation. Sensory attenuation is the reduction of the precision afforded to self-generated sensory stimuli, intuitively exemplified as the inability to tickle oneself (Blakemore et al., 2000; Shergill et al. 2003). Impairments in sensory attenuation have long been implicated in theoretical models of hallucinations and delusions (Fletcher and Frith, 2009; Frith and Done, 1989) – as well as other phenomena in schizophrenia, such as improved performance in the force-matching paradigm (Shergill et al., 2005) and a loss of N1 suppression during speech (Ford et al., 2007). We did not include them in our model because failures of sensory attenuation in schizophrenia may best be described as occurring at lower levels of the sensorimotor hierarchy (Shergill et al., 2005; Cassidy et al., 2018), whereas our model deals with very high-level functions – the recognition of linguistic content. We therefore felt that including failure of low-level sensory attenuation would overcomplicate the model by mixing mechanisms operating at different hierarchical levels. As such our model is unlikely to capture certain aspects of AVH phenomenology that are thought to be related to failure of sensory attenuation, such as the attribution of external agency to voices, or the phenomenon of thought insertion (Sterzer et al., 2016). However, our generative model does allow us to examine what mechanisms operate at higher levels, where content is inferred, in isolation.

## CONCLUSION AND FUTURE DIRECTIONS

In this paper we have demonstrated simulations of agents experiencing hallucinations that are or are not consistent with the environmental context. On a Bayesian (active inference) reading of perception, when likelihood precision was reduced, agents experienced hallucinations congruent with the environmental context. When we additionally induced anomalous prior beliefs about trajectories (i.e., the sequence of words in a narrative), the agent experienced out-of-context hallucinations. These out-of-context hallucinations were reminiscent of disordered thought in psychotic subjects, suggesting that there may be common mechanisms underlying hallucinations and thought disorder. This work suggests that we may distinguish between two, possibly interconnected, aspects of computational psychopathology. First, the genesis of hallucinatory percepts depends upon an imbalance between prior beliefs and sensory evidence. This idea generalises across psychotic hallucinations, and the induction of false positive inferences in healthy subjects used in empirical studies. Second, disordered content associated with a false positive inference may depend upon additional pathology that affects prior beliefs about the trajectories of hidden states. Beliefs about unusual transitions may be a more latent trait in schizophrenia, which become fully manifest only when the precision of sensory evidence is compromised.

## Acknowledgements

DB is supported by a Richard and Edith Strauss Fellowship (McGill University) and the Fonds de Recherche du Quebec-Santé (FRQS). TP is supported by the Rosetrees Trust (Award Number 173346). RAA was supported by an NIHR University College London Hospitals Biomedical Research Centre Postgraduate Fellowship and is now an MRC Skills Development Fellow (MR/S007806/1). KJF is a Wellcome Principal Research Fellow (Ref: 088130/Z/09/Z).

## Software note

All simulations were run using the DEM toolbox included in the SPM12 package. This is open-access software provided and maintained by the Wellcome Centre for Human Neuroimaging and can be accessed here: http://www.fil.ion.ucl.ac.uk/spm/software/spm12/

## Footnotes

1 When the outcomes are sensory samples, the likelihood precision plays a role of a sensory precision in predictive coding formulations.

